# Ecological niche modelling of the wattled crane (*Bugeranus carunculatus*) suggest range expansion and contraction during the Pleistocene

**DOI:** 10.1101/406710

**Authors:** Daniel Augusta Zacarias

**Affiliations:** Universidade Eduardo Mondlane, Escola Superior de Hotelaria e Turismo de Inhambane. Cidade de Inhambane, Bairro Chalambe-2, Av. Samora Machel, Caixa Postal 75

**Keywords:** climate change, range shifts, disjunct distribution, paleodistribution modelling, birds, range expansion-contraction theory

## Abstract

This paper attempts to understand the potential effects of historical climate changes on the distribution of the wattled crane (*Bugeranus carunculatus*). The potential distribution in past and present climatic scenarios is investigated through ensemble distribution modelling of 232 independent and sparsely distributed occurrence records. Potential effects of climate change were evaluated by means on niche overlap and niche gains and losses across time scales. Massive range expansion was observed from the Last Interglacial (LIG) era to the Last Glacial Maximum (LGM), with loss of suitability in most areas of western Africa and an increase in suitability across southern and eastern Africa. From the LGM, climate suitability tended to establish in southern and eastern Africa with slight disjunction in the mid-Holocene, a trend that was maintained through current distribution. Results indicate the presence of southern and northern refugia, with massive range expansion in central populations. These results support the idea that the current disjunct distribution of the wattled crane is driven by climate oscillations during the Pleistocene that generated range expansion and retraction of the species and also support the hypothesis that the current occurrence of the species is driven by other factors such as food and habitat availability.

## 1. Introduction

Disjunct distribution of species has always intrigued biogeography (Deng et al., 2015; Beatty and Provan, 2013). However, consensus do exist that species distribution is determined by the interaction between biotic, abiotic and historical factors (Peterson et al., 2011; Brown et al. 1996). While this interaction is commonly understood when attempting to explain the distribution of continuous species, the situation gets rather complex when attempting to explain complex disjunct distributions. Several approaches and studies have been developed to understand why single species are located in different areas, with two traditional hypotheses having received extensive consideration: vicariance and dispersal (González et al., 2014; Lozano-Jaramillo et al., 2014).

On one side, the vicariance hypothesis, assumes that current disjunct distributions are relicts of former continuous distributions (Karanth, 2003), caused by range contractions due to changes in climatic conditions that might have affected habitat suitability (Cox and Moore, 2005). As such, species disjunct distribution is a result of the fragmentation of ancestral wide distribution caused by the appearance of any barrier. On the other side, the dispersal hypothesis assumes that disjunct distributions are a result of long-distance dispersal to suitable habitats (Cox and Moore, 2005) through a pre-existing dispersal barrier (Kropf et al., 2006). These patterns are extensively studied through species paleodistribution modelling (Wethey et al., 2016; Pena et al., 2014), phylogenetic analysis of divergence (Deng et al., 2015; Hawlitschek et al., 2015) or a combination of both (Thesing et al., 2016; Vitorino et al., 2016; Lozano-Jaramilo et al., 2014). These hypotheses, though, are not mutually exclusive and consensus is that long-term maintenance of disjunct distributions is dependent on the environmental unsuitability of surrounding areas and/or adaptation to different environmental conditions of geographically distinct areas (Lozano-Jaramilo et al., 2014; Gatson, 2003).

In times of increasing biodiversity loss, disjunct species distribution can be a great challenge to species long-term persistence (Byrne et al., 2016; Rubinoff et al., 2015; Marchant et al., 2015), mainly because land degradation and fragmentation can heavily affect smaller and discontinuous ranges (Sandel et al., 2011) accelerating species extinction (Sobral-Souza et al., 2015; Harris and Pimm, 2008). In addition, disjunct distributions challenge biodiversity conservation, as conservation needs to understand and choose whether to protect the species as a whole or as special management units (Measey and Tolley, 2011). This is particularly difficult when historical information is lacking and conservation actions need to be urgently considered (Pena et al., 2014; Measey and Tolley, 2011).

One of the key features of cranes worldwide is the dominance of disjunct distributions. Of the 15 extant crane species (Duan and Fuerst, 2001; Meine and Archibald, 1996), only the sandhill crane, *Grus canadensis*, appears to have a single continuous distribution. The wattled crane, *Bugeranus carunculatus*, presents an intriguing disjunct distribution across southern and eastern Africa consisting of three major populations, namely: the south-central population along the Okavango delta and the Kafue flats, the South African population and the Ethiopian populations (Bento et al., 2007; Jones et al., 2006; Meine and Archibald, 1996). Like the majority of crane species, the wattled crane is threatened with extinction, being classified as vulnerable (VU; A2acde+3cde; C1+2a) by the International Union for Conservation of Nature (IUCN) and enlisted at the Appendix II of the Convention on International Trade in Endangered Species of Wild Fauna and Flora (CITES) and the Convention on Migratory Species (CMS).

Little is known about the reasons behind the origin of the current disjunction of *B. carunculatus* distribution, with existing knowledge considering that south African and Ethiopian populations are relicts of a past continuous distribution (Beilfuss et al., 2003). However, considering the climatic oscillations during the Pleistocene period, it is plausible to assume that the sequence of relatively warmer and wetter (Last Interglacial – LIG) and cooler and drier (Last Glacial Maximum – LGM) climate conditions (Gavashelishvili and Tarkhnishvili, 2016) might have driven changes in the geographic distribution of species, associated to the contraction of lowland forest and savannah and the ability of the species to track suitable climatic conditions.

In addition, considering the relative dependence of the species in relation to lowland floodplain habitats and the presence of *Eleocharis spp*., it is also plausible to postulate that the disjunct distribution of *B. carunculatus* might also be a result of its climate specialist behaviour that has maintained its distribution with respect to climate and vegetation types. This paper explores these assumptions, based on distribution modelling of the species through the Afrotropical realm in 4 consecutive climatic timescales: the Last Interglacial (LIG), the Last Glacial Maximum (LGM), the Mid-Holocene (Mid-H) and contemporary climate.

## 2. Material and methods

### 2.1 Species occurrence records

Occurrence records were retrieved from the Global Biodiversity Information Facility (GBIF; GBIF.org), encompassing historical (n = 47) and contemporary records (n = 6224). After checking for data inconsistencies, duplicates and errors, 3625 records were retained for further analysis (Table S1). These records were then spatially filtered (Boria et al., 2013) considering a species home range of 16 km^2^ (McCann and Benn, 2006), resulting in 232 independent records (Figure 1) that were then used to model the potential climatic distribution of the species. Spatial filtering was done using the SDMtoolbox (Brown et al., 2017; Brown, 2014;) in ArcGIS 10.5.

**Fig 1.**
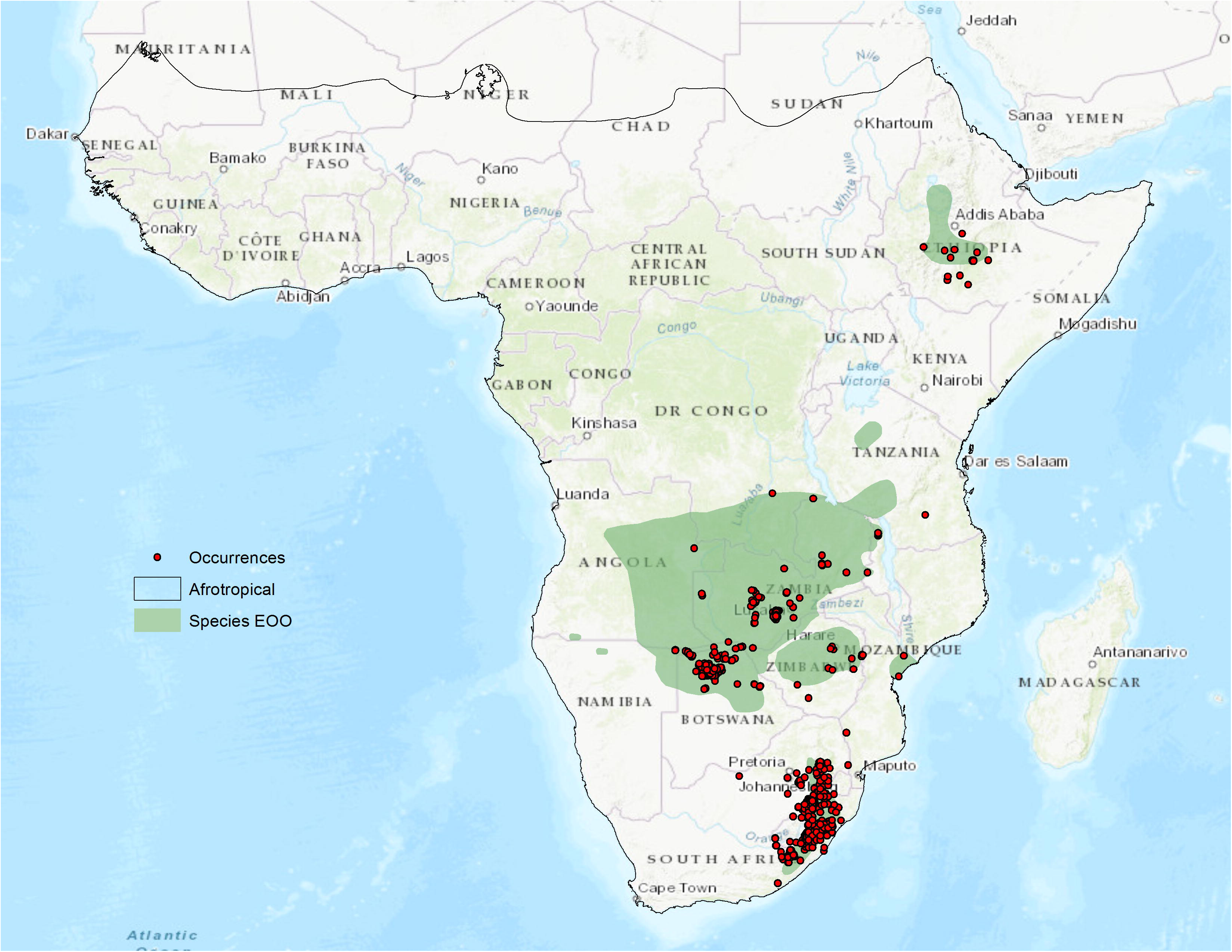
The extent of occurrence (EOO) of the wattled crane in Africa and the occurrence records (n = 232) used for distribution modelling, obtained after data cleaning and filtering. Note that the number of occurrence records available at the GBIF database (Table S1) is much larger that the number displayed here. For colours, the reader is referred to the web version of the paper.

### 2.2 Bioclimatic variables

Current bioclimatic variables associated to annual mean temperature (bio1), temperature seasonality (bio4), maximum temperature of the warmest period (bio5), annual precipitation (bio12), precipitation seasonality (bio15) and precipitation of the driest quarter (bio17) were downloaded from the World Climate Database (http://www.worldclim.org; Hijmans et al., 2005) at a spatial resolution of 2.5 min.

These variables were chosen because they (i) have been extensively applied in studies of birds’ response to climate change in several environments (Ribeiro et al., 2016; Barbet-Massin and Jetz 2015; Reside et al., 2012) and (ii) fulfil some of the most important biological requirements for birds (Bateman et al., 2016; Reside et al., 2012). For past climatic conditions, the same bioclimatic variables were used, at the same resolution. As for the Mid-Holocene (Mid-H; ∼ 6kya) and Last Glacial Maximum (LGM; ∼ 22kya), climate variables spanned three global ocean circulation models (AOGCMs), namely: CCSM4, MIROC-ESM and MPI-ESM-P. For the Last Inter-Glacial (LIG; ∼120kya – 140kya), data were downloaded at a resolution of 30 arc-seconds and rescaled to the same resolution as other variables (2.5 min) using the rescale toolbox of the SDMtoolbox (Brown et al. 2017; Brown 2014) in ArcGIS.

### 2.3 Species distribution modelling

Potential climatic distribution of the species was predicted using *biomod2* (Thuiller et al., 2016) an ensemble platform for species distribution modelling. Climatic suitability was first predicted for current climate scenario and then projected into past scenarios, using nine algorithms (Table S2). For each model, occurrence records were randomly partitioned into training (75%) and test (25%) datasets, with 5 replications. The True Skills Statistics (TSS) and Area Under the Curve (AUC) evaluated models predictive capability. For each climate scenario (except for the LIG, were only dataset was available) an ensemble of all nine algorithms was calculated (Tinker et al., 2015; Loyola et al., 2013), resulting in consensus maps of potential climatic distribution. As for the LGM and Mid-H, maps of potential climate suitability across AOGCMs were averaged to generate a map of potential suitability for each climate scenario per climatic era (Hovick et al., 2015; Loyola et al., 2013).

### 2.4 Potential distribution, range shifts and refugia

Maps resulting from species distribution modelling indicate the climatic suitability of the species across a continuous gradient ranging from 0 to 1, were 0.5 indicates random distribution and 1 optimum distribution. To understand areas with higher probability of species presence, all maps were converted into a binary distribution with values equal or lower that 0.5 indicating areas with no presence of the species and otherwise indicating presence of the species. The resulting binary maps were used to assess potential range shifts across timescales and gains and losses of climatic suitability. For every pair of climatic timescales the earlier prediction was subtracted by the older prediction to generate range variations of −1 (range contraction) and 1 (range expansion).

For each timescale, the number of occupied grids was counted to calculate the size of habitat contraction, expansion and stability. All maps were summed to calculate potential *refugia* areas, i.e. areas that were potentially suitable for the species since the Last Interglacial climatic scenario (Ban et al. 2016; Keppel et al. 2011). Niche similarity (or dissimilarity) across timescales was calculated based on the Schoener’s *D* (Schoener, 1968) and Warren’s I statistics (Warren et al., 2009) using the *nicheOverlap* function of the *dismo* package (Hijmans et al., 2016) in R (R Core Team, 2017). These statistics compute the degree of ecological overlap between two datasets at a scale of 0 to 1, with values near 0 associated to niche dissimilarity (no overlap) between two datasets and values near 1 associated to high niche similarity (niche overlap) (Broennimann et al., 2012).

## 3. Results

Models evaluation indicated that, overall, all models had predictive capacity better than random (Table S2). The lower TSS and AUC values were associated to the Surface Range Envelop (0.710 ± 0.054 and 0.855 ± 0.027, respectively) and the highest was associated to the Random Forest algorithm (0.902 ± 0.023 and 0.980 ± 0.006, respectively). The potential distribution of the species through time is displayed in Figure 2, with the predicted current distribution largely matching the current distribution as estimated by the IUCN extent of occurrence, suggesting equilibrium with current climate. As for the past, the distribution of the wattled crane demonstrated distribution during the LGM and progressive shrinkage in the following scenarios, suggesting climatic instability during the Pleistocene (Figure 2; Figure S1).

**Fig 2.**
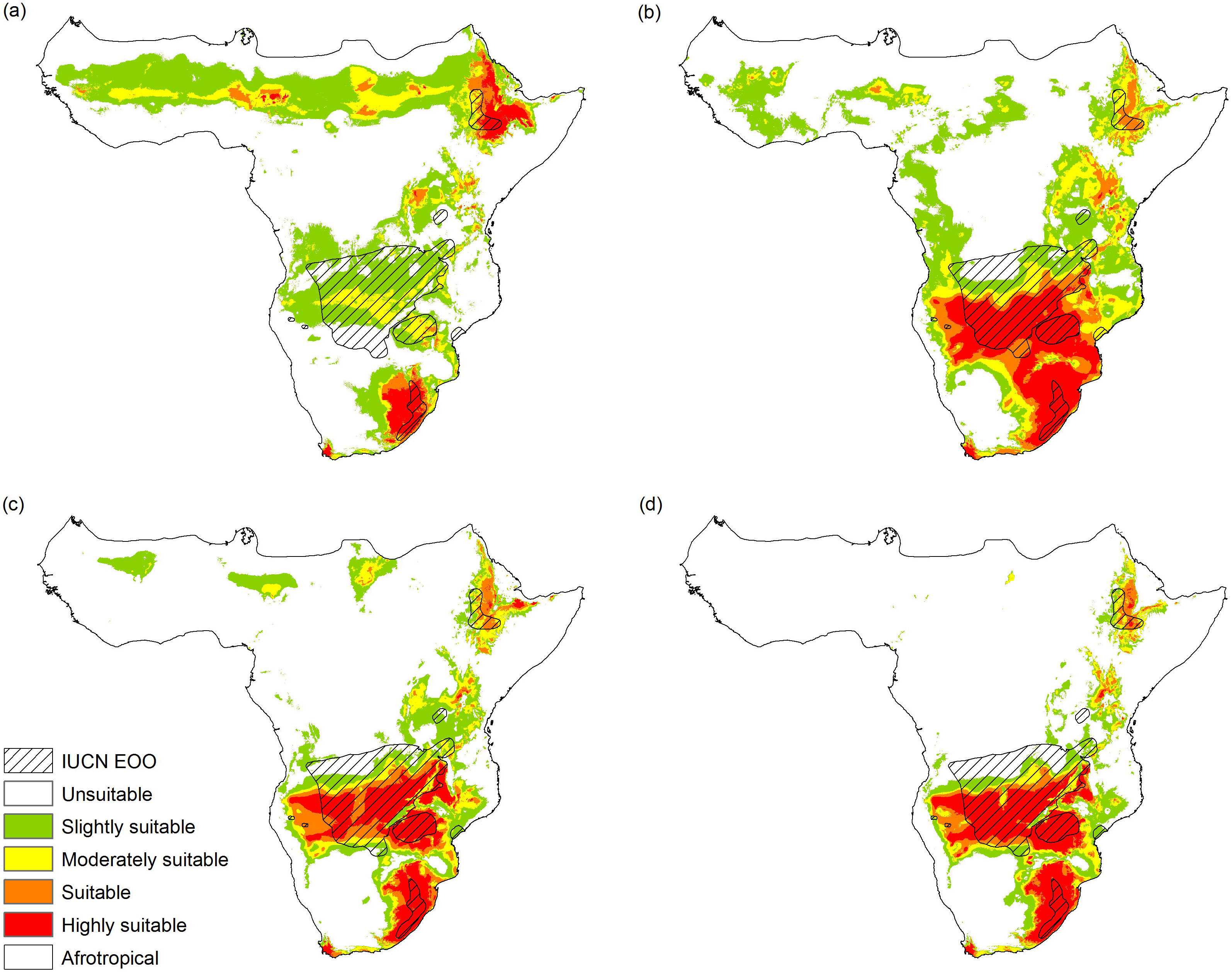
Predicted suitability for the wattled crane, *Bugeranus carunculatus*, across the Afrotropical realm. A) Last Interglacial (∼130 kya); B) Last Glacial Maximum (∼21kya); C) Mid-Holocene (∼6ka); D) Current climate (0kya). Warmer colours correspond to areas with higher probability of occurrence. Black line represents the borders of the Afrotropical realm (Olson et al. 2001) and dashed lines represent the IUCN EOO (IUCN, 2017). For colours, the reader is referred to the web version of the paper.

A high dissimilarity was found between suitable climatic areas between the LIG and LGM (Schoener’s D = 0.255, Warren’s I = 0.398, Table 1) and similarity between the suitable areas in Mid-H and current climatic scenarios (Schoener’s D = 0.860, Warren’s I = 0.895), suggesting that there has been an improvement of the climatic conditions for the species from the LGM to the mid-Holocene and relative stability in the last 6000 years. These results suggest a dynamic distribution of climatically suitable areas for *B. carunculatus*, highlighting South African and Ethiopian populations as possible refugia (Figure 3).

**Fig 3.**
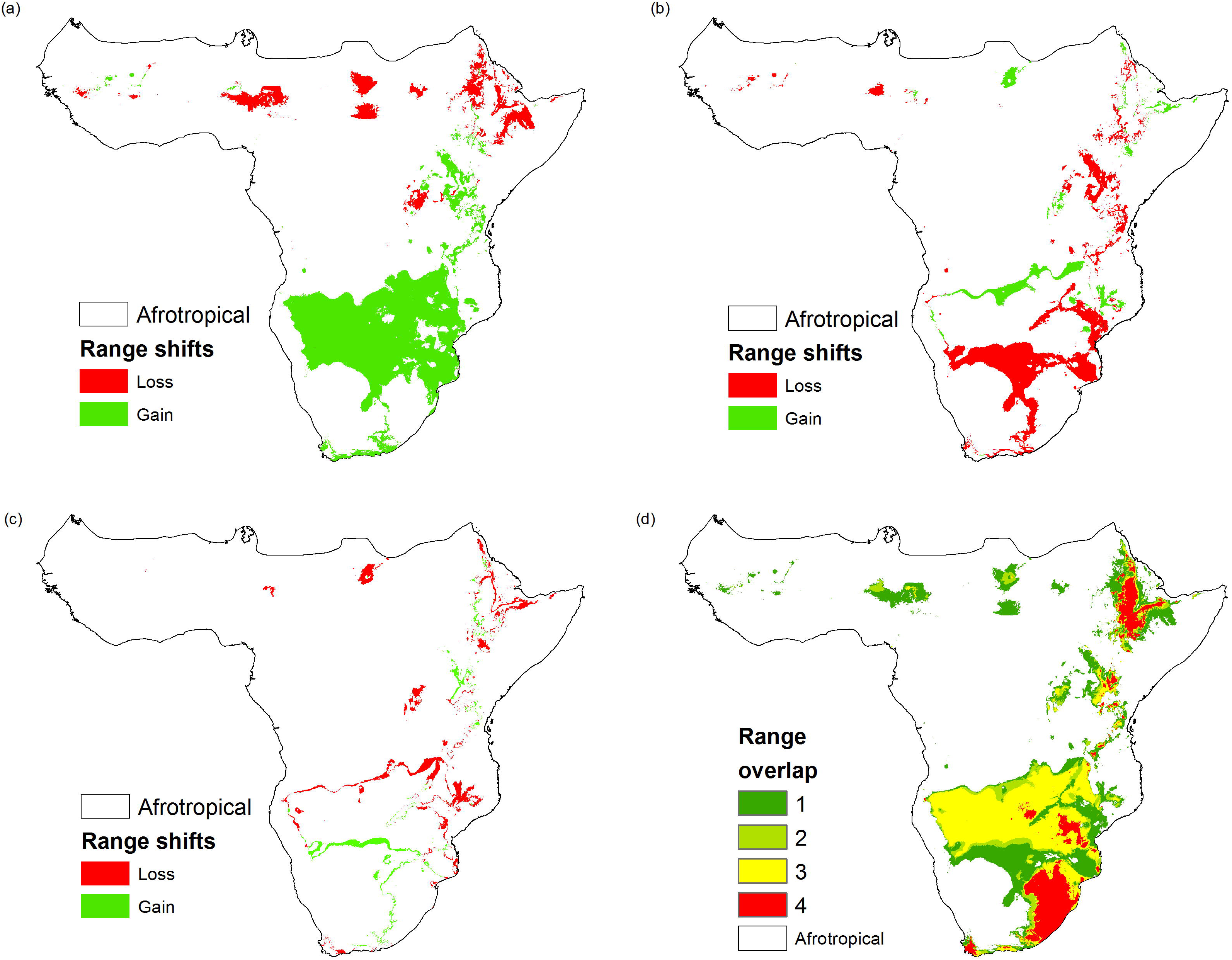
Patterns of climatic range shifts for the wattled cranes in the Afrotropical realm through time. A) Range shifts between the LIG and LGM; B) Range shifts between the LGM and Mid-H; C) Range shifts between the Mid-H and present climate scenarios; D) Overlap of climate suitability across all time slices. In A-C, red indicate areas that lost suitability and green indicates areas that gained suitability. In D) areas in red represent potential climatic *refugia* for the species, or areas that have maintained highly suitable through all timescales. For colours, the reader is referred to the web version of the paper.

**Table 1.** Patterns of range shifts and niche overlap between timescales. Columns 3-5 represent the number of cells predicted as gained (G) or lost (L) across two time slices and the corresponding balance (GL). Columns 6-7 represent measures of niche similarity and dissimilarity (Schoener’s *D* and Warren’s *I*) across time scales.

| From | To | G | L | GL | $D$ | $I$ |
| --- | --- | --- | --- | --- | --- | --- |
| ~ 130kya | ~ 22kya | 146,449 | 30,753 | 115,896 | 0.255 | 0.398 |
| ~ 22kya | ~ 6kya | 17,430 | 63,143 | -45,713 | 0.679 | 0.775 |
| ~ 6kya | 0k | 9590 | 21,073 | -11,483 | 0.860 | 0.895 |

## 4. Discussion

Results herein reported, suggest that at any period in the past the wattled crane had scattered distribution across the Afrotropics, but benefited from an improvement of bioclimatic conditions during the transition from the LIG to the LGM that increased suitability in southern, the area that is currently occupied by the core population of the species. The modelling process accurately matched the current distribution and highlighted the presence of two refugia in eastern and southern Africa, thus providing evidence for the glacial refugia hypothesis (Brito, 2005).

The increase of suitability in southern Africa, though, is a complex process with complex understanding, but can be associated to the biogeography of the present-day central Africa and the dynamics of the Congo-Zambezi system that influenced the distribution and evolution of tropical Africa’s biota (Schultheiß et al., 2014) due to the formation of the East African Rift System (EARS). This complex system affected the distribution of tropical flora and fauna, but had a decisive role in creating, disrupting, redirecting and connecting freshwater systems (Schultheiß et al., 2014; Roberts et al., 2012) essential to the survival of the wattled crane. This connectivity between the Congo and the Zambezi river system can explain the persistence of wattled cranes in southern Africa, due to the prevalence of extensive shallow floodplains that provide sufficient habitat and resources for the species.

Results, herein, contradict the former assumption that South African and Ethiopian populations are relicts of a former broader continuous distribution of wattled cranes in Africa (Beilfuss et al. 2003), but rather establish that these populations might be climatic refugia of the species that have maintained effective numbers through time. In addition, the current distribution of the species appears to be also a result of low dispersal ability of the species (Jones et al. 2006). *B. carunculatus* current distributions appear to follow lowlands surrounding the major afromontane ecosystems in Africa, that might imply that the rise of the Rift Valley ecoregion might have been responsible in separating the Ethiopian population from the south-central population. However, reasons behind possible separation between south-central population and the South African population remain unclear, although the limited dispersal hypothesis might be considered the starting point.

Nevertheless, results herein, support the hypothesis that wattled cranes has once in time established across western Africa (Birdlife International, 2018). As illustrated in Figure 2, there have been areas with scattered climatic suitability through time that could have possibly harboured cranes. Due to climate oscillations during the Quaternary, associated to the deepening of major swell in Western Africa and consequently disappearance of shallow floodplains (Miller and Gosling, 2014). In addition, the distribution of the wattled cranes can also be attributed to main climatic events during early years of the quaternary that created conditions for the emergence of barriers to the dispersal of the species (Burke and Wilkinson, 2016; Mark and Osmaston, 2008) mainly by the lack of important food resources (Jones et al. 2006). Adding to this, it has been emphasized that the climatic regime associated to the last glaciations during the LGM have created conditions for the prevalence of swells and shallow floodplains nearby the afromontane complexes (Burke and Wilkinson, 2016; Osmaston and Harrison, 2005) that might have created sufficient conditions for the persistence of the species due to its high adaptability to new environments.

Although this study does not integrate phylogenetic analysis, its results when associated to the biology of the species provide sufficient evidence to support the hypothesis that the current distribution of *B. carunculatus* is a result of complex range-shifts during the Quaternary that improved suitable conditions for the core population in southern Africa and enabled the persistence of two refugia for the south African and Ethiopian populations. Nevertheless, the incorporation of phylogenetic studies might shed new lights into this complex debate and provide additional insights into species long-term persistence and guide management practices. Previous attempts to estimate genetic relatedness between south-central and South African populations has indicated that South African populations have comparable levels of genetic variation as to the south-central, despite lack of gene flow between them as a result of lack of matrilineal gene flow across the region (Jones et al. 2006).

This study is the first experience to attempt to understand the origin of the current disjunct distribution of the wattled crane. Results indicated that climate changes during the Quaternary drove the current distribution of the species by improving environmental conditions in southern Africa and isolating two refugia, one in South Africa and the other in Ethiopia. As such, results support the hypothesis that the current distribution of *B. carunculatus* is a vicariance-based process associated to the great adaptability potential of the species to survive in areas that are currently not suitable but, still, provide enough food resources such as the Zambezi delta in central Mozambique. Further work on the intra and interspecific genetic variation and phylogenetic analysis of extant and populations is recommended to further uncover the colonization history and to identify whether the predicted climatic refugia are, simultaneously, genetic refugia and to validate the hypothesis of no dispersal among *B. carunculatus* populations.

Nevertheless, the present distribution indicate that relatively high proximity to heavy occupied settlements that can hinder future prospects of species survival, especially when considering the increasing demand for water supply that has resulted in extensive drainage of water systems. Considering that this species is highly sensitive to human disturbance (Morrison and Bothma, 1998), successful conservation is a challenging process that will require balancing surrounding land-uses and reduce disturbance of the species.

## 5. Acknowledgements

I wish to thank the GBIF team and contributors for making their data freely available. Software and package developers are also acknowledged. This paper benefited from earlier discussion with Leticia Braga. This research did not receive any specific grant from funding agencies in the public, commercial or not-for-profit sectors.

## List of tables

Table S1. *Bugeranus carunculatus* presence records at the Global Biodiversity Information Facility.

Table S2. Models performance

Table S3. Differences (paired *t*-test) of climatic suitability across timescales

## Figure captions

FigS1. Predicted a) Last Interglacial (∼130 kya), b) Last Glacial Maximum (∼21 kya), c) Mid-Holocene (∼6 kya) and d) current (0 kya) distribution of the wattled crane. Superimposed is the IUCN extent of occurrence (EOO).

